# Active Bed Microclimate Regulation Modulates Sleep Architecture, Autonomic, and Central Nervous System Activity during Sleep

**DOI:** 10.64898/2026.09.07.749920

**Authors:** Gary Garcia-Molina, Megha Rajam Rao, Trevor Winger, Pavlo Chernega, Yehor Shcherbakov, Erwin Veneros, Kathryn Jean Reid, Daniela Grimaldi, Eus Van Someren, Phyllis Zee

## Abstract

Skin temperature and bed microclimate are established modulators of human sleep, but most prior work has relied on passive thermal materials. We used a smart bed capable of heating or cooling the microclimate to investigate how six temperature settings affect sleep architecture, autonomic activity, and electroencephalographic (EEG) activity. Eighteen adults (9 male, 9 female; mean age 44.5 years) completed two overnight polysomnography sessions with bed temperature assigned via Latin-square counterbalancing across three-hour segments of high cooling, medium cooling, low cooling, off, low heating, and medium heating. Mixed-effects regression models evaluated manipulation checks on induced thermal changes, the probability of sleep versus wake, and the probability of each sleep stage (N1, N2, N3, REM), while secondary analyses assessed sleep deepening, heart rate, and EEG spectral power. Temperature manipulations significantly affected sleep and physiological outcomes. All five active settings shifted sleep toward greater depth, increasing the odds of slow-wave sleep (odds ratios 1.21 to 3.65, largest for high cooling) and reducing the odds of light non-rapid eye movement sleep relative to off. Heating accelerated early sleep deepening relative to cooling without significantly altering sleep onset latency, whereas cooling produced a graded reduction in heart rate. EEG analyses revealed distinct cortical signatures for heating and cooling conditions. These findings demonstrate that active bed microclimate regulation modulates sleep architecture, autonomic activity, and cortical activity during sleep. The nonlinear pattern of effects suggests thermal “sweet spots” that may vary across the night and individuals, supporting the development of individualized programmable temperature-regulation strategies for sleep optimization.

## Introduction

Sleepiness and vigilance are closely associated with core body (CBT) and skin temperature (ST). Within the thermoneutral range, high CBT and low ST promote vigilance, while low CBT and high ST facilitate sleep. Sleep initiation is associated with the circadian rhythms of CBT and skin temperature, with habitual sleep onset occurring during the CBT decline and concomitant ST increase (Van Someren, 2006).

The nocturnal CBT decline is facilitated by peripheral vasodilation which increases cutaneous blood flow. The resulting increase in skin temperature (Zoccoli, Silvani and Franzini, 2013). facilitates heat dissipation into the micro-environment. The rise in skin temperature is particularly evident during the transition from wakefulness to sleep (Kräuchi et al., 1999). Under the controlled conditions of a constant routine protocol, the distal-to-proximal skin temperature gradient (DPG) was identified as the strongest physiological predictor of sleep onset latency, outperforming core body temperature, heart rate, melatonin, and subjective sleepiness ratings (Kräuchi et al., 1999, 2000). Critically, this relationship was deemed functional rather than merely correlative, as diverse thermoregulatory interventions consistently demonstrated that greater pre-sleep distal vasodilation corresponded with shorter time to sleep onset.

A causal role for skin warming in shortening sleep onset latency was first demonstrated experimentally by Raymann, Swaab and Van Someren (2005), who used a water-perfused thermosuit to independently manipulate proximal (trunk and limbs) and distal (hands and feet) skin temperature. They found that warming of the proximal skin by less than 1°C reduced sleep onset latency by approximately 27% (at least 3 minutes), while distal manipulation alone was less effective, possibly due to the relatively narrow temperature range applied. A subsequent study by the same group (Raymann, Swaab and Van Someren, 2008) extended these findings to sleep depth and architecture, confirming that skin temperature feeds back on sleep-regulating systems in the brain across both young adults and elderly participants.

More recent work focused on promoting heat loss. Herberger et al. (2024) investigated the impact of sleeping on a high heat capacity mattress (HHCM) on sleep in a multi-center study that involved 72 individuals of varying age, gender, and body-mass index. HHCMs promote body cooling by facilitating heat transfer from the body’s core to the mattress. The main findings were that sleeping on a HHCM increased slow wave sleep by 7.5 (SD 21.6) minutes and decreased heart rate by 2.36 (SD 1.08) beats/min. Reid et al. (2021) reported benefits of sleeping on a HHCM in a group of 24 post-menopausal women (aged 40 to 75) in a single-blind, counterbalanced, crossover design. Sleeping on the HHCM selectively increased slow-wave sleep and slow-oscillation activity during the first non-rapid eye movement/rapid eye movement sleep cycle only.

These passive-mattress studies establish that passive facilitation of heat loss from the body can promote slow-wave sleep and reduce heart rate. It would be interesting to investigate whether effects of active heating and cooling as previously demonstrated using a thermosuit can also be accomplished using a more feasible and user-friendly thermal manipulation. Leveraging the ability of the Sleep Number Climate 360 bed (Sleep Number, Minneapolis, MN) to actively control the temperature of the microclimate in either direction, a study was conducted to test the effect of temperature on sleep architecture, cardiorespiratory activity, and electroencephalogram-derived metrics during sleep.

## Methods

### Participants

Eighteen volunteers (9 male, 9 female) were recruited from Sleep Number customers, with mean age 44.5 years (SD: 6.7) and mean BMI 29.0 kg/m² (SD: 4.4). All participants owned a Climate 360 smart bed and resided within easy reach of the Sleep Health Center of Northwestern University in Chicago, IL, USA. The study was approved by the Institutional Review Board of Northwestern University (protocol STU00217800). Written informed consent was obtained from all participants.

### Study design

Each participant completed two overnight laboratory sleep sessions, experiencing different temperature programs across nights. Each night consisted of three consecutive three-hour segments, with a different temperature setting applied to each segment (see Figure 1). Temperature programs were designed to cover six settings across two nights: high cooling (HC), medium cooling (MC), low cooling (LC), Off, low heating (LH), and medium heating (MH). Assignment of settings to segments was counterbalanced across participants using a Latin-square design, ensuring that each setting appeared equally often across segment positions and was not confounded with time of night. Bedtimes and the corresponding temperature program onset were individualized to each participant’s self-reported typical bedtime to maximize ecological validity

**Figure 1.**
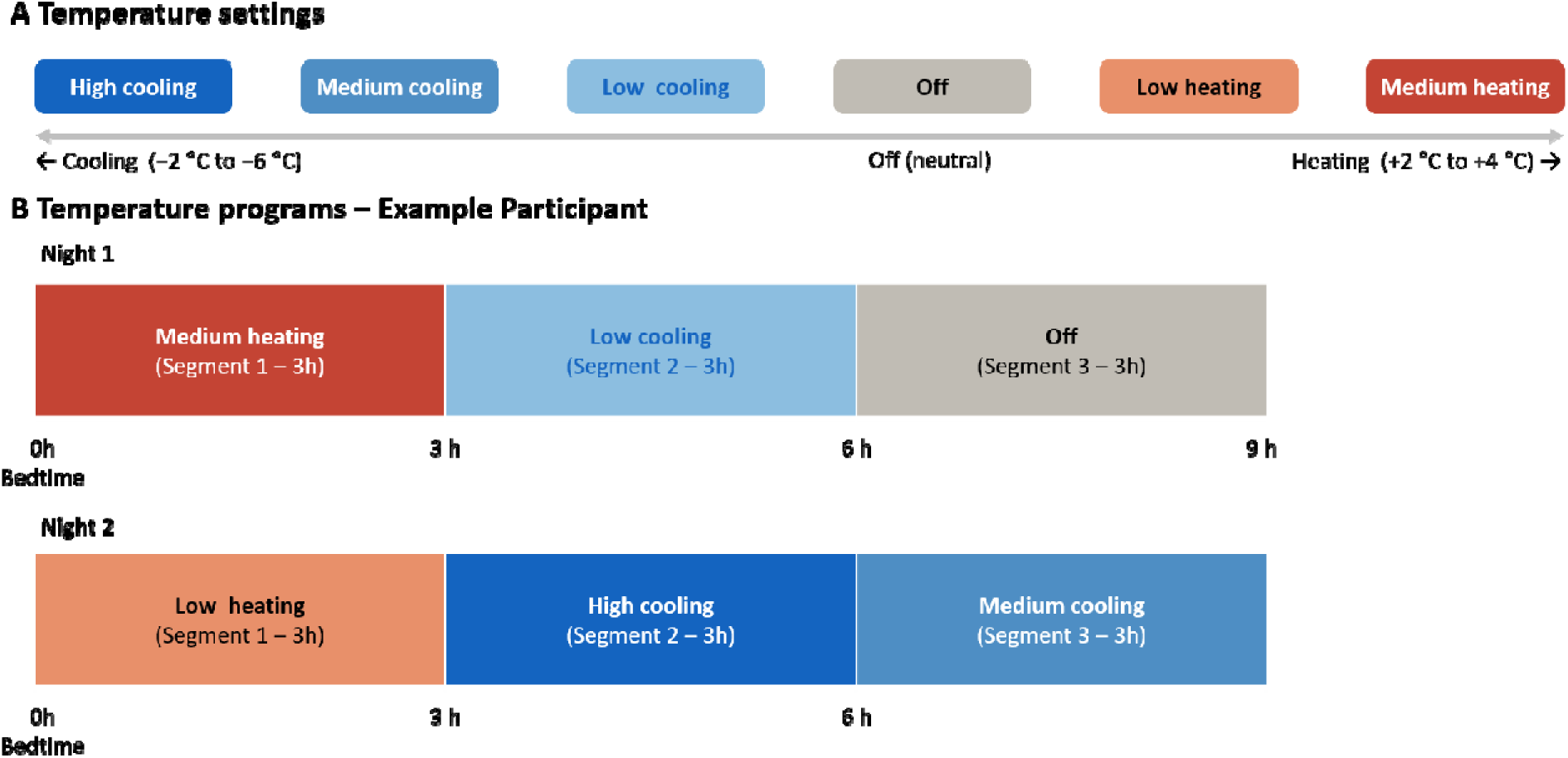
Study design. Panel A: the six temperature settings available to the bed system, spanning active cooling (−2°C to −6°C) to active heating (+2°C to +4°C). Panel B: the two-night temperature program for Participant 2, illustrating the Latin-square counterbalanced assignment of settings to segments. Each night comprised three consecutive three-hour segments, each set to a different temperature. Across the two nights, each participant received all six settings exactly once, with the order counterbalanced across participants to avoid confounding setting with segment position or night.

### Climate 360 Smart Bed System

The Climate 360 bed modulates the temperature of the enclosed space (the microclimate) between the sleeper’s body, bedding, and mattress surface. An integrated thermal module provides independent, side-specific control by circulating conditioned air within an internal mattress layer that exchanges heat with the microclimate through the mattress surface. In heating mode, external air is warmed to 30°C (low heating) or 32°C (medium heating); the resulting microclimate temperature increases by approximately +2°C and +4°C, respectively. In cooling mode, heat absorbed in the internal mattress layer is removed and exhausted to the room environment, decreasing microclimate temperature by approximately 2°C, 4°C, or 6°C for low, medium, or high cooling settings, respectively. Because thermal exchange occurs through the mattress surface rather than direct air injection into the space between the sleeper and bedding, the intervention does not disrupt mattress comfort or require external ducting.

An array of five temperature sensors arranged along a strip from the bed edge toward the center sampled microclimate temperature at ∼0.5 Hz (Figure 2). For analyses, sensor T1 (nearest to the bed edge) most consistently tracked temperature-setting changes unperturbed by body coverage variability and was therefore used to verify intervention delivery (manipulation check).

**Figure 2.**
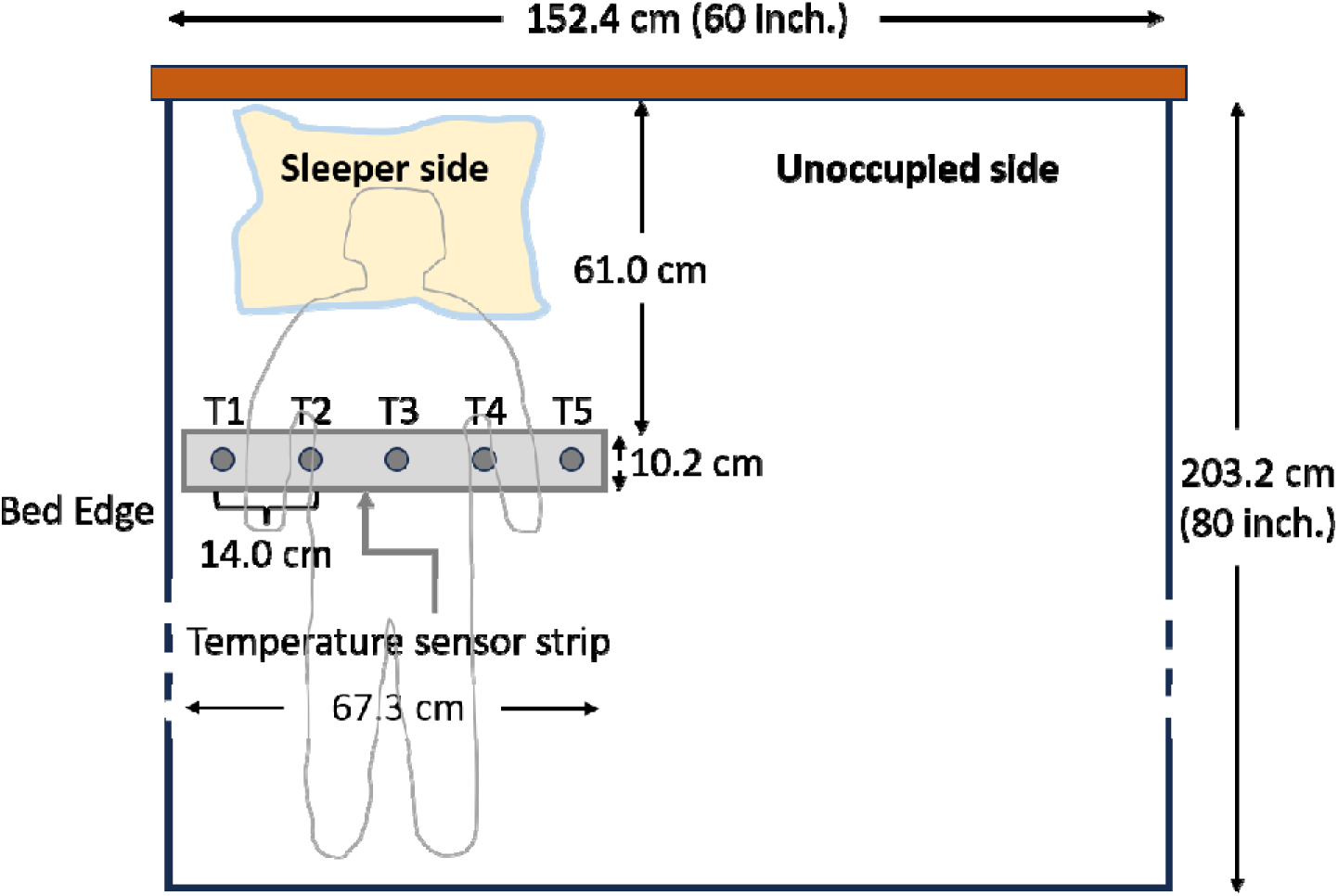
Location of the temperature sensor strip on the Climate 360 smart bed (queen size, 152.4 × 203.2 cm). The strip was positioned on the participant’s sleeping side, with sensor T1 nearest the bed edge and sensors spaced 14.0 cm apart (total strip length 67.3 cm). Data from the sleeping-side strip were used in all analyses; no sensor strip was recorded on the unoccupied side.

### Physiological measurements

#### Polysomnography

Standard polysomnography signals (Neurofax EEG-1100 Digital EEG Acquisition System, Nihon-Kohden, Tokyo, Japan) were recorded at 200 Hz, including electroencephalography (EEG) from locations F3, F4, C3, C4, O1, O2, electromyography (EMG), electro-oculography (EOG), electrocardiography (ECG), respiratory signals, and limb movement electrodes. Electrode impedances were reduced during application and maintained at < 5 kOhms for EEG/EOG and < 10 kOhms for EMG, in accordance with standard PSG technical guidance (American Academy of Sleep Medicine, 2023). Sleep stages were manually scored in 30-second epochs according to American Academy of Sleep Medicine (AASM) criteria (American Academy of Sleep Medicine, 2023) by an experienced sleep scientist who was blinded to the temperature program during scoring.

#### Skin temperature

Distal skin temperature (DST) was sampled with 0.0625°C resolution using an iButton (DS1922L, Maxim Integrated) positioned on the right foot arch. Proximal skin temperature (PST) was measured with an iButton positioned below the left clavicle. The distal-to-proximal gradient (DPG) was calculated as DST minus PST.

#### Heart rate

ECG signals recorded during polysomnography were processed to detect R-peaks using a wavelet-based algorithm (Goldberger et al., 2000; Moody and Mark, 2001). Heart rate (HR) was computed for each 30-second epoch.

#### EEG spectral analysis

EEG signals were analyzed in the spectral domain to compute mean power in Delta (0.5–4 Hz), Theta (4–8 Hz), Alpha (8–12 Hz), Sigma/Spindle (11–16 Hz), and Beta (15–30 Hz) bands for each 30-second epoch. Spectral power in each band was normalized relative to total broadband power within the same epoch using the formula: normalized band power (dB) = 10 × log₁₀(band power / total power); log refers to log_10_ throughout. This normalization expresses each frequency band as a proportion of total EEG activity within that epoch, removing epoch-to-epoch and between-participant differences in overall signal amplitude.

The logarithm of the ratio of Delta to Beta power was calculated as an indicator of sleep deepening during the early sleep period (Gorgoni et al., 2021); higher log(Delta/Beta) values reflect deeper sleep. Centering of all post-sleep-onset signals (delta/beta, heart rate, and skin temperature) was performed relative to the epoch immediately preceding sleep onset, defined as the first epoch scored as sleep; this serves as a conventional clinical anchor point within the gradual transition from wakefulness to sleep, rather than a sharp physiological boundary. Epoch-wise statistical comparisons of these post-onset dynamics were performed at 30-second resolution to identify epochs of significant difference between temperature conditions (uncorrected p<0.05).

### Statistical analysis

#### Outcome structure

Analyses were organized into three tiers. First, a manipulation check was run to confirm that temperature settings produced the intended changes in bed microclimate temperature and skin temperature. Second, the primary outcome characterized the effect of temperature setting on sleep architecture, evaluated first as the probability of being asleep versus awake, and second as the probability of being in each sleep stage (N1, N2, N3, REM). Third, secondary outcomes characterized the dynamics of falling asleep (sleep onset latency and the early post-onset trajectory of delta/beta ratio, heart rate, and skin temperature) and the effect of temperature setting on heart rate and EEG spectral power across the full night, stratified by sleep stage.

#### Mixed-effects modeling

Mixed-effects linear and logistic regression models accommodated the hierarchical data structure, with 30-second epochs nested within nights nested within subjects. Time since sleep onset was included as a second-order covariate to account for the natural nonlinear progression of sleep and sleep stage probabilities, as well as EEG and ECG features changing across the night.

Effects of temperature settings were initially intended to us dummy coding, where at any given epoch at most one of the variables high cooling (HC), medium cooling (MC), low cooling (LC), low heating (LH), and medium heating (MH) could have the value 1, and all others the value 0. Off was chosen as the reference condition, in which case HC, MC, LC, LH, and MH all had the value 0. Data inspection however showed that bed temperature changed only slowly after each switch of temperature setting. To obtain adequate regressors of bed temperature acting on sleep, the slow increase and decay curves were estimated using physics-informed exponentially increasing and decreasing functions, convolved with binary indicators of setting status. The resulting variables (HC, MC, LC, LH, MH) slowly increased from 0 to attain the value 1 only after approximately 30 minutes following a switch to that setting and slowly decreased from 1 to attain the value 0 only after approximately 45 minutes following a switch away from that setting. The previous and current setting dummy values add up to 1 at any time point, except when the previous or current setting is Off, in which case the value is between 0 and 1. All reported results use these slowly changing dummy variables rather than dummy variables that change unrealistically immediately with a switch in temperature setting.

#### Use of aggregated cooling and heating conditions

The manipulation check and primary outcome analyses are reported exclusively at the level of the five individual temperature settings, since aggregating across the three cooling and two heating levels would obscure dose-dependent effects that are themselves of interest at this stage. For two of the secondary, post-sleep-onset point-wise analyses (delta/beta and skin temperature dynamics), settings were additionally aggregated into Cooling, Off, and Heating groups to increase statistical power for epoch-wise testing. We note that this aggregation pools an unequal number of intensity levels (three cooling levels versus two heating levels) and is therefore reported as a power-recovery strategy for hypothesis-generating, exploratory comparisons rather than as a primary or confirmatory analysis; this imbalance is revisited as a limitation in the Discussion.

## Results

### Manipulation check

#### Bed temperature

Bed temperature strip sensors T1 and, to a slightly lesser extent, T2, positioned from the bed edge toward the center, reliably tracked the monotonic relationship between temperature settings and the resulting thermal changes. Sensors located closer to the bed center may be less reflective of the programmed setting due to local heat exchange with the sleeper’s body and direct contact with bedding. For sensor T1, temperatures changed by −2.11°C (high cooling), −1.62°C (medium cooling), −0.90°C (low cooling), +3.39°C (low heating), and +4.67°C (medium heating) relative to off (Table 1). All values in Table 1 are statistically significant: corresponding confidence intervals do not include zero.

**Table 1.** Estimated regression coefficients (95% confidence intervals) for each bed temperature sensor (T1–T5) across microclimate settings, relative to Off.

|  | T1 | T2 | T3 | T4 | T5 |
| --- | --- | --- | --- | --- | --- |
| **High Cooling** | $-2.11^{\circ}\text{C}$<br>[$-2.20, -2.01$] | $-1.70^{\circ}\text{C}$<br>[$-1.82, -1.58$] | $1.01^{\circ}\text{C}$<br>[$0.90, 1.12$] | $2.51^{\circ}\text{C}$<br>[$2.42, 2.60$] | $1.64^{\circ}\text{C}$<br>[$1.53, 1.75$] |
| **Medium Cooling** | $-1.62^{\circ}\text{C}$<br>[$-1.71, -1.52$] | $-2.02^{\circ}\text{C}$<br>[$-2.13, -1.90$] | $1.67^{\circ}\text{C}$<br>[$1.56, 1.78$] | $2.29^{\circ}\text{C}$<br>[$2.20, 2.38$] | $1.27^{\circ}\text{C}$<br>[$1.16, 1.38$] |
| **Low Cooling** | $-0.90^{\circ}\text{C}$<br>[$-0.99, -0.80$] | $-0.88^{\circ}\text{C}$<br>[$-0.99, -0.76$] | $1.92^{\circ}\text{C}$<br>[$1.81, 2.04$] | $2.56^{\circ}\text{C}$<br>[$2.47, 2.64$] | $1.83^{\circ}\text{C}$<br>[$1.72, 1.95$] |
| **Low Heating** | $3.39^{\circ}\text{C}$<br>[$3.30, 3.49$] | $2.14^{\circ}\text{C}$<br>[$2.03, 2.26$] | $1.69^{\circ}\text{C}$<br>[$1.58, 1.80$] | $0.59^{\circ}\text{C}$<br>[$0.50, 0.68$] | $0.16^{\circ}\text{C}$<br>[$0.05, 0.27$] |
| **Medium Heating** | $4.67^{\circ}\text{C}$<br>[$4.58, 4.76$] | $3.26^{\circ}\text{C}$<br>[$3.14, 3.38$] | $2.00^{\circ}\text{C}$<br>[$1.89, 2.12$] | $1.47^{\circ}\text{C}$<br>[$1.38, 1.55$] | $2.77^{\circ}\text{C}$<br>[$2.66, 2.88$] |

Model: $T_{\text{sensor}} \sim \text{HC} + \text{MC} + \text{LC} + \text{LH} + \text{MH} + (1 \mid \text{participant})$ . Mixed-effects linear regression (with participant and night random intercepts) modeled temperature settings as gradual, continuous effects over time (relative to Off baseline), estimating how each (fully) active setting changes sensor temperature. The model was fitted using restricted maximum likelihood.

#### Skin temperature

DST was 33.8 ± 1.8 °C on average across all Off observations. DST responded broadly in line with manipulation intensity for cooling conditions, decreasing by −1.01°C (high cooling) and −1.25°C (medium cooling) relative to off, but increasing slightly by +0.39°C under low cooling. Both heating conditions produced comparable increases in DST of +1.28°C (low heating) and +1.25°C (medium heating).

PST was 34.4 ± 1.3 °C on average across all Off observations. PST responded to manipulation with a more complex pattern: high cooling produced a modest PST increase of +0.42°C, while medium cooling and low cooling both decreased PST, by −0.90°C and −0.84°C respectively. Low heating increased PST by +0.52°C, whereas medium heating reduced it by −0.41°C.

The DPG was −0.57 ± 2.2 °C on average across all Off observations. DPG responses reflected the combined distal and proximal dynamics: high cooling reduced DPG by −1.53°C, whereas medium cooling produced a smaller reduction of −0.47°C. Low cooling increased the DPG by +1.22°C. Both heating conditions produced positive changes in DPG shifts with a moderate increase (+0.68°C) by low heating and a large increase by medium heating (+1.63°C). Full regression coefficients and 95% confidence intervals are reported in Table 2.

**Table 2.** Estimated regression coefficients (95% confidence intervals) for distal (DST), proximal (PST), and distal-to-proximal gradient (DPG) skin temperature under each microclimate setting, relative to Off.

|  | Distal | Proximal | DPG |
| --- | --- | --- | --- |
| **High Cooling** | $-1.01^{\circ}\text{C}$<br>[$-1.07, -0.95$] | $0.42^{\circ}\text{C}$<br>[$0.35, 0.48$] | $-1.53^{\circ}\text{C}$<br>[$-1.63, -1.44$] |
| **Medium Cooling** | $-1.25^{\circ}\text{C}$<br>[$-1.32, -1.19$] | $-0.90^{\circ}\text{C}$<br>[$-0.97, -0.83$] | $-0.47^{\circ}\text{C}$<br>[$-0.57, -0.37$] |
| **Low Cooling** | $0.39^{\circ}\text{C}$<br>[$0.33, 0.46$] | $-0.84^{\circ}\text{C}$<br>[$-0.91, -0.77$] | $1.22^{\circ}\text{C}$<br>[$1.12, 1.32$] |
| **Low Heating** | $1.28^{\circ}\text{C}$<br>[$1.22, 1.34$] | $0.52^{\circ}\text{C}$<br>[$0.46, 0.59$] | $0.68^{\circ}\text{C}$<br>[$0.58, 0.77$] |
| **Medium Heating** | $1.25^{\circ}\text{C}$<br>[$1.18, 1.31$] | $-0.41^{\circ}\text{C}$<br>[$-0.48, -0.34$] | $1.63^{\circ}\text{C}$<br>[$1.53, 1.73$] |

Model: $T_{\text{skin}} \sim \text{HC} + \text{MC} + \text{LC} + \text{LH} + \text{MH} + (1 \mid \text{participant})$ . Mixed-effects linear regression (with participant-specific random intercepts) modelled temperature settings as gradual, continuous effects over time (relative to Off baseline), estimating how each (fully) active setting changes skin temperature. The model was fitted using restricted maximum likelihood.

### Primary outcome: effect of temperature setting on sleep architecture

Separate mixed-effects logistic regression models were fitted to estimate temperature setting effects (all settings versus Off) on the probability of Sleep (N1+N2+N3+REM) versus Wake, and of each individual stage (N1, N2, N3, REM, and Wake) versus any other stage. Defining each stage as a binary outcome (that stage versus all others) decreases the compositional dependency that would otherwise complicate a joint analysis of all stages together. Thus, each model is statistically independent and the standard odds ratio interpretation is preserved. The model structure included participant-level random intercepts and regressors for second-order changes in stage probability across the night. Results are presented in Table 3.

**Table 3.** Odds ratios (95% confidence intervals) for the probability of being asleep and for the probability of being in each sleep stage, across temperature settings relative to Off. * p < 0.05 vs Off.

| Setting | Asleep | N1 | N2 | N3 | REM | Wake |
| --- | --- | --- | --- | --- | --- | --- |
| High-Cooling | 1.06<br>[0.91, 1.24] | **0.77\*** [0.64, 0.94] | **0.74\*** [0.67, 0.81] | **3.65\*** [3.08, 4.33] | **1.20\*** [1.05, 1.37] | 0.94<br>[0.81, 1.09] |
| Medium-Cooling | **1.46\*** [1.24, 1.71] | **0.75\*** [0.60, 0.93] | 1.00<br>[0.90, 1.11] | **1.21\*** [1.01, 1.45] | **1.43\*** [1.24, 1.65] | **0.69\*** [0.58, 0.81] |
| Low-Cooling | **2.84\*** [2.37, 3.40] | **0.72\*** [0.58, 0.90] | **1.16\*** [1.04, 1.30] | **1.38\*** [1.13, 1.68] | **1.44\*** [1.24, 1.67] | **0.35\*** [0.29, 0.42] |
| Low-Heating | **1.83\*** [1.53, 2.17] | **0.67\*** [0.55, 0.83] | 1.09<br>[0.98, 1.21] | **1.77\*** [1.48, 2.11] | **1.18\*** [1.03, 1.37] | **0.55\*** [0.46, 0.65] |
| Medium-Heating | **3.21\*** [2.69, 3.83] | **0.72\*** [0.58, 0.88] | 1.04<br>[0.94, 1.15] | **2.64\*** [2.20, 3.17] | 1.10<br>[0.96, 1.26] | **0.31\*** [0.26, 0.37] |

Boldened text indicates $p < 0.05$ .

#### Sleep versus wake

The overall probability of being asleep (second column in Table 3) was significantly increased relative to Off under medium cooling (OR = 1.46), low cooling (OR = 2.84), low heating (OR = 1.83), and medium heating (OR = 3.21). High cooling was the exception, with no significant change in the probability of being asleep (OR = 1.06, p > 0.05). This pattern points to a non-linear relationship in which moderate thermal stimulation in either direction promotes sleep maintenance, while the most intense cooling setting does not.

#### Sleep stage architecture

N1 probability was significantly reduced relative to Off across all five active temperature conditions (ORs: 0.67–0.77), indicating a consistent shift away from light NREM sleep regardless of the direction or magnitude of the temperature manipulation. N3 probability was significantly increased under all five conditions (ORs: 1.21–3.65), with the largest effect observed for high cooling (OR = 3.65 [3.08, 4.33]), followed by medium heating (OR = 2.64 [2.20, 3.17]). Wake probability was significantly reduced under four of the five conditions, with the largest reductions observed for medium heating (OR = 0.31 [0.26, 0.37]) and low cooling (OR = 0.35 [0.29, 0.42]); the exception was high cooling, which did not significantly alter wake probability (OR = 0.94 [0.81, 1.09]).

REM probability was significantly increased under four conditions (ORs: 1.18–1.44), with no significant effect under medium heating (OR = 1.10 [0.96, 1.26]). N2 showed the most heterogeneous pattern across conditions: high cooling significantly reduced N2 probability (OR = 0.74 [0.67, 0.81]), low cooling significantly increased it (OR = 1.16 [1.04, 1.30]), and the remaining three conditions produced no significant change.

Taken together, all active temperature conditions shifted sleep architecture away from N1 and toward deeper NREM sleep. The dissociation observed for high cooling, namely the largest gain in slow-wave sleep of any condition but no corresponding reduction in wake probability or increase in the overall probability of being asleep, is an interesting architectural finding which is examined further in the Discussion alongside the EEG spectral findings in Table 4, which suggest a possible underlying mechanism.

**Table 4.** Normalized EEG spectral power (dB, 95% confidence intervals) during sleep epochs under each active temperature condition, relative to Off. * p < 0.05 vs Off (two-tailed).

| Setting | Delta (dB) | Theta (dB) | Alpha (dB) | Sigma (dB) | Beta (dB) |
| --- | --- | --- | --- | --- | --- |
| High Cooling | -0.03<br>[-0.29, 0.24] | **-0.46\*** [-0.58, -0.33] | **-0.61\*** [-0.73, -0.49] | **-0.30\*** [-0.44, -0.17] | **-1.25\*** [-1.41, -1.10] |
| Medium Cooling | 0.21<br>[-0.07, 0.49] | -0.03<br>[-0.16, 0.10] | -0.00<br>[-0.13, 0.13] | **0.30\*** [0.16, 0.45] | **-0.24\*** [-0.41, -0.08] |
| Low Cooling | **0.46\*** [0.16, 0.76] | -0.06<br>[-0.20, 0.08] | **-0.37\*** [-0.51, -0.24] | **0.42\*** [0.27, 0.57] | -0.10<br>[-0.27, 0.07] |
| Low Heating | **0.94\*** [0.66, 1.22] | -0.13<br>[-0.26, 0.01] | **-0.29\*** [-0.42, -0.16] | -0.05<br>[-0.20, 0.09] | **-0.57\*** [-0.73, -0.41] |
| Medium Heating | **0.47\*** [0.20, 0.74] | **-0.15\*** [-0.28, -0.02] | **-0.38\*** [-0.51, -0.26] | -0.07<br>[-0.21, 0.07] | **-0.78\*** [-0.94, -0.62] |

Model: normalized band power (dB) ~ HC + MC + LC + LH + MH + time + time<sup>2</sup> + (1 | session). Mixed-effects linear regression (with random intercepts per recording session) modelled temperature settings as gradual, continuous effects over time (relative to Off baseline), with a second-order time-of-night covariate to account for the natural drift in EEG spectral composition across the night. Boldened text indicates $p < 0.05$ .

#### EEG spectral power as a mechanistic correlate of architecture effects

To clarify the cortical activity underlying the sleep-architecture effects shown in Table 3, mixed-effects linear regression models restricted to sleep-scored epochs, accounting for sleep stage and time of night, were used to evaluate normalized EEG spectral power across frequency bands (Table 4). These analyses are presented here, immediately following the architecture results, because they offer a possible mechanistic explanation of the largest N3 gain associated with high cooling.

High cooling produced a pattern qualitatively distinct from all other settings: broad suppression of activity in frequency bands other than delta, with significant reductions in theta (−0.46 dB), alpha (−0.61 dB), sigma (−0.30 dB), and beta (−1.25 dB) relative to Off.

Heating conditions showed contrasting spectral signature: both low heating and medium heating significantly increased delta power (+0.94 dB and +0.47 dB, respectively) while concurrently suppressing alpha and beta power, indicating enhanced slow-wave activity together with reduced higher-frequency cortical engagement. Low cooling occupied an intermediate position, significantly increasing both delta (+0.46 dB) and sigma (+0.42 dB) power while reducing alpha (−0.37 dB). Medium cooling produced a more modest shift, selectively increasing sigma power (+0.30 dB) and attenuating beta (−0.24 dB).

Across all five active conditions, beta power was reduced or unchanged relative to Off, indicating that active thermal modulation in either direction trends to attenuates high frequency cortical activity.

### Secondary outcomes

Secondary outcomes evaluated the sleep onset process, as well as additional autonomic and cortical effects of temperature settings that are not captured by the primary sleep architecture outcomes presented in the previous section.

Temperature settings were considered individually and were also aggregated into Cooling, Off, and Heating groups to enhance statistical power.

#### Sleep onset latency

There was no significant effect of the initial temperature program setting on sleep onset latency (SOL), neither with individual regressors for each active setting versus Off (Figure 3A; linear mixed-effects model, likelihood ratio test: χ²(5) = 4.81, p = 0.440); nor with active settings aggregated into cooling and heating versus Off (Figure 3B; χ²(2) = 1.34, p = 0.512).

**Figure 3.**
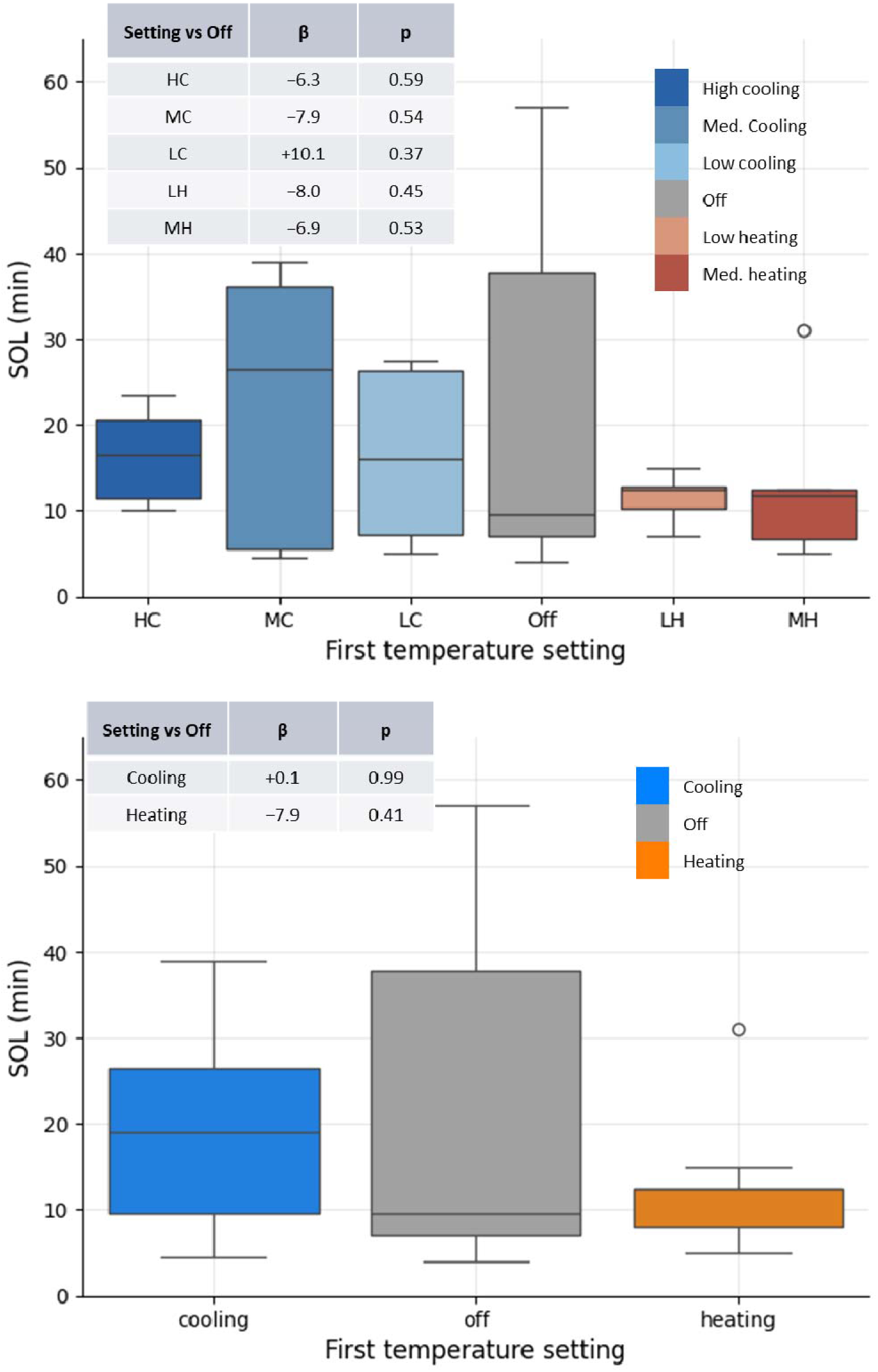
Sleep onset latency versus first temperature setting. (A) All temperature settings and intensities thereof. (B) Aggregated into heating and cooling regardless of intensity.

#### Delta/beta ratio dynamics following sleep onset

The epoch-by-epoch delta/beta ratio development over time was evaluated with the epoch immediately preceding sleep onset as baseline. Epoch-wise two-sided Mann-Whitney U tests were applied at each epoch to identify of difference between conditions (uncorrected p < 0.05).

When examined across individual temperature settings (Figure 4A), delta/beta ratio trajectories diverged across the first hour following sleep onset. Heating conditions (low heating and medium heating) showed the most rapid increase, reaching the highest delta/beta values within the first 60 minutes. Cooling conditions (high cooling, medium cooling, low cooling) and Off showed comparatively attenuated deepening trajectories.

**Figure 4.**
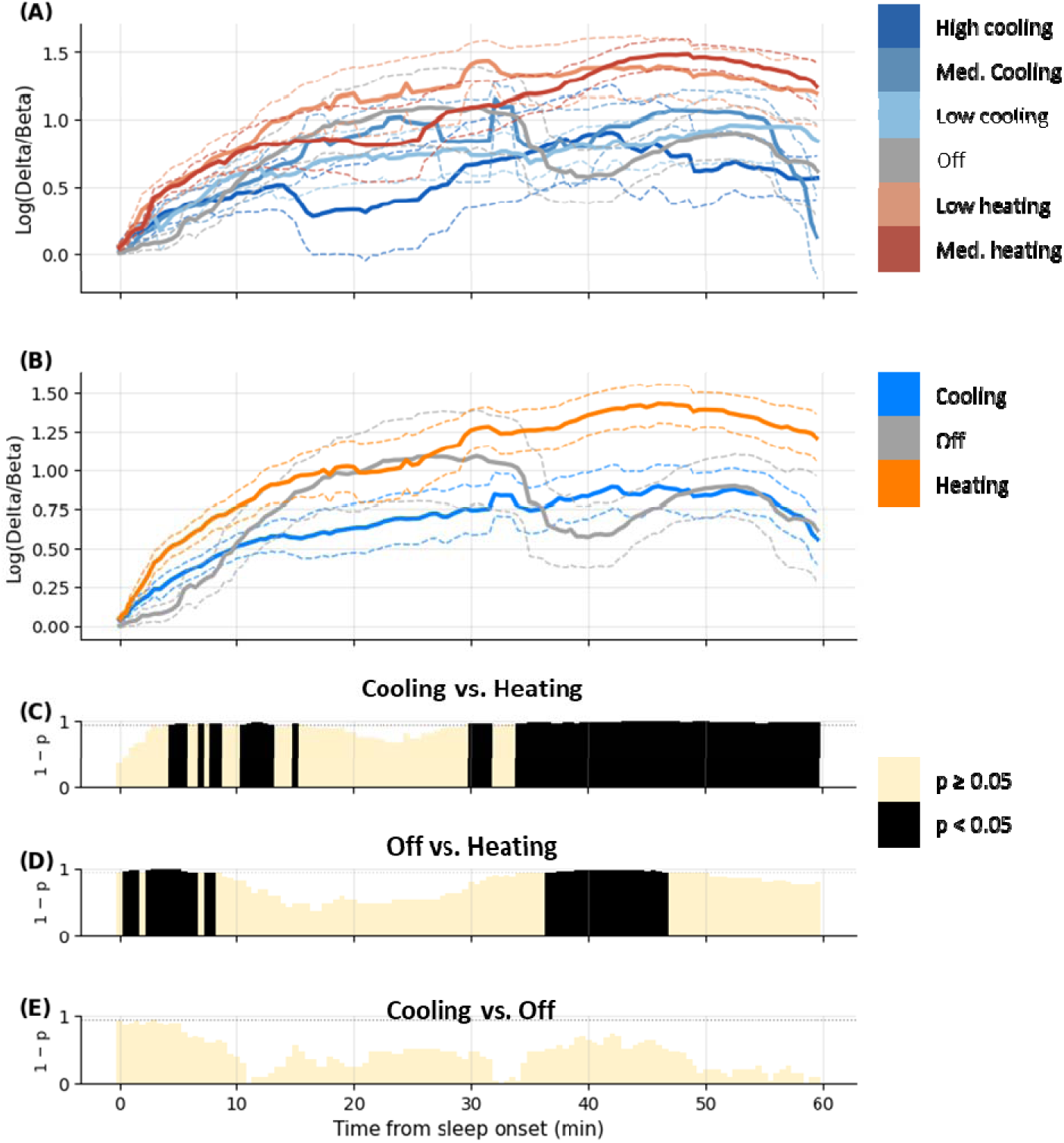
Delta/beta ratio dynamics from sleep onset as a function of the first temperature setting. (A) Mean log(Delta/Beta) for each individual setting; dashed lines represent ±1 standard error. (B) Aggregated into Cooling, Off, Heating groups. (C–E) Point-wise statistical significance (1 − p) from two-sided Mann-Whitney U tests at 30-second resolution. Black bars: p < 0.05; tan bars: p ≥ 0.05. (C) Cooling vs Heating. (D) Off vs Heating. (E) Cooling vs Off.

When settings were collapsed into Cooling, Off, and Heating groups (Figure 4B), these differences became clearer. Heating produced greater sleep deepening than cooling starting approximately 30 minutes after sleep onset, with significant epochs from ∼30 minutes onward through the remainder of the monitored period (Figure 4C; Mann-Whitney U, p < 0.05). Heating also produced significantly greater sleep deepening than Off during the first ∼10 minutes after sleep onset and again from approximately 35 to 45 minutes (Figure 4D; Mann-Whitney U, p < 0.05). No point-wise differences with p < 0.05 were found between cooling and Off at any epoch (Figure 4E; Mann-Whitney U, p ≥ 0.05 throughout).

#### Heart rate

Mixed-effects linear regression models adjusting for sleep stage and time of night demonstrated significant stage-dependent effects of temperature manipulation on heart rate relative to the Off condition (Table 5). At the aggregate level, cooling produced a graded reduction in heart rate, with effect magnitude increasing monotonically with cooling intensity (high cooling: −3.06 bpm; medium cooling: −2.10 bpm; low cooling: −1.18 bpm). Whereas low heating induced only a small but statistically significant reduction (−0.28 bpm), medium heating resulted in an increase (+1.74 bpm).

**Table 5.** Regression coefficients (bpm, 95% confidence intervals) for the effect of temperature setting on heart rate by sleep stage, relative to Off. * p < 0.05 vs Off.

| Setting | Overall | N1 | N2 | N3 | REM | Wake |
| --- | --- | --- | --- | --- | --- | --- |
| High-Cooling | **−3.06\***<br>[−3.28, −2.84] | **−4.38\***<br>[−5.29, −3.47] | **−5.15\***<br>[−5.46, −4.85] | **−2.92\***<br>[−3.45, −2.38] | **−3.21\***<br>[−3.74, −2.68] | −0.13<br>[−0.96, 0.71] |
| Medium-Cooling | **−2.10\***<br>[−2.33, −1.86] | **−2.76\***<br>[−3.81, −1.70] | **−3.45\***<br>[−3.76, −3.14] | +0.04<br>[−0.56, 0.64] | **−2.39\***<br>[−2.91, −1.87] | **−1.67\***<br>[−2.56, −0.79] |
| Low-Cooling | **−1.18\***<br>[−1.43, −0.93] | **−2.61\***<br>[−3.66, −1.55] | **−1.96\***<br>[−2.29, −1.62] | **+0.99\*** [0.32, 1.66] | **−2.12\***<br>[−2.63, −1.62] | **−1.49\***<br>[−2.49, −0.49] |
| Low-Heating | **−0.28\***<br>[−0.52, −0.05] | **−1.33\***<br>[−2.31, −0.35] | **−1.14\***<br>[−1.45, −0.82] | −0.30<br>[−0.94, 0.34] | **−0.82\***<br>[−1.34, −0.30] | −0.38<br>[−1.43, 0.67] |
| Medium-Heating | **+1.74\*** [1.51, 1.98] | +1.01<br>[−0.01, 2.04] | +0.30<br>[−0.01, 0.61] | **+3.89\*** [3.25, 4.54] | **+1.92\*** [1.42, 2.41] | **+4.47\*** [3.47, 5.47] |

Model: $HR \sim HC + MC + LC + LH + MH + SleepStage + time + time^2 + (1 | session)$ . Mixed-effects linear regression (with random intercepts per recording session) modelled temperature settings as gradual, continuous effects over time (relative to Off baseline). Sleep stage (N1, N2, N3, REM; reference: Wake) and a second-order time-of-night covariate were included to account for the natural variation in heart rate across stages and across the night. Per-stage columns are from separate models restricted to epochs of that stage, without the sleep-stage covariate. Off served as the reference condition. Boldened values indicate $p < 0.05$ .

Stage-specific analyses revealed heterogeneity underlying these overall effects. Cooling-related reductions were most pronounced during lighter and intermediate NREM sleep (N1 and N2), where all cooling levels significantly decreased heart rate, with the largest absolute effect in N2 under high cooling (−5.15 bpm). Responses during deep sleep (N3) deviated from this pattern and were non-monotonic: high cooling remained associated with a significant reduction (−2.92 bpm), medium cooling showed no measurable effect, and low cooling increased heart rate (+0.99 bpm). REM sleep, by contrast, exhibited consistent and significant heart rate reductions across all cooling conditions (approximately −2 to −3 bpm).

Heating conditions displayed a distinct pattern. The overall increase in heart rate under medium heating was driven primarily by large and significant elevations in N3 (+3.89 bpm), REM (+1.92 bpm), and especially wake (+4.47 bpm), while effects in lighter sleep stages (N1 and N2) were small and not statistically significant. Low heating, by contrast, produced modest decreases in heart rate limited to N1, N2, and REM, with no detectable effects in N3 or wake.

In the immediate post-sleep-onset window, point-wise comparisons of heart rate showed minimal statistically significant differences between conditions, despite a descriptive upward drift in cooling conditions (see Supplementary Figure S4). This is consistent with the limited power of uncorrected point-wise testing in this sample and is superseded by the stage-adjusted all-night model above.

#### Distal-to-proximal skin temperature gradient during sleep onset

The distal-to-proximal gradient was evaluated from sleep onset using the epoch immediately preceding sleep onset as baseline. DPG is presented as the headline skin-temperature secondary outcome because it summarizes the joint distal and proximal dynamics in a single physiologically interpretable measure.

DPG trajectories diverged by temperature setting over the first 60 minutes following sleep onset (Figure 5). Cooling conditions drove DPG into increasingly negative territory, reaching approximately −1°C below baseline, while low heating and Off drove a sustained positive shift of approximately +1°C; medium heating produced a comparatively flat trajectory. Cooling produced significantly lower DPG than both Off and heating across much of the monitored period (Mann-Whitney U, p < 0.05); no significant point-wise differences were observed between Off and heating at any epoch. Full point-wise statistical breakdowns by condition pair, along with individual DST and PST trajectories, are reported in Supplementary Figures S1–S3.

**Figure 5.**
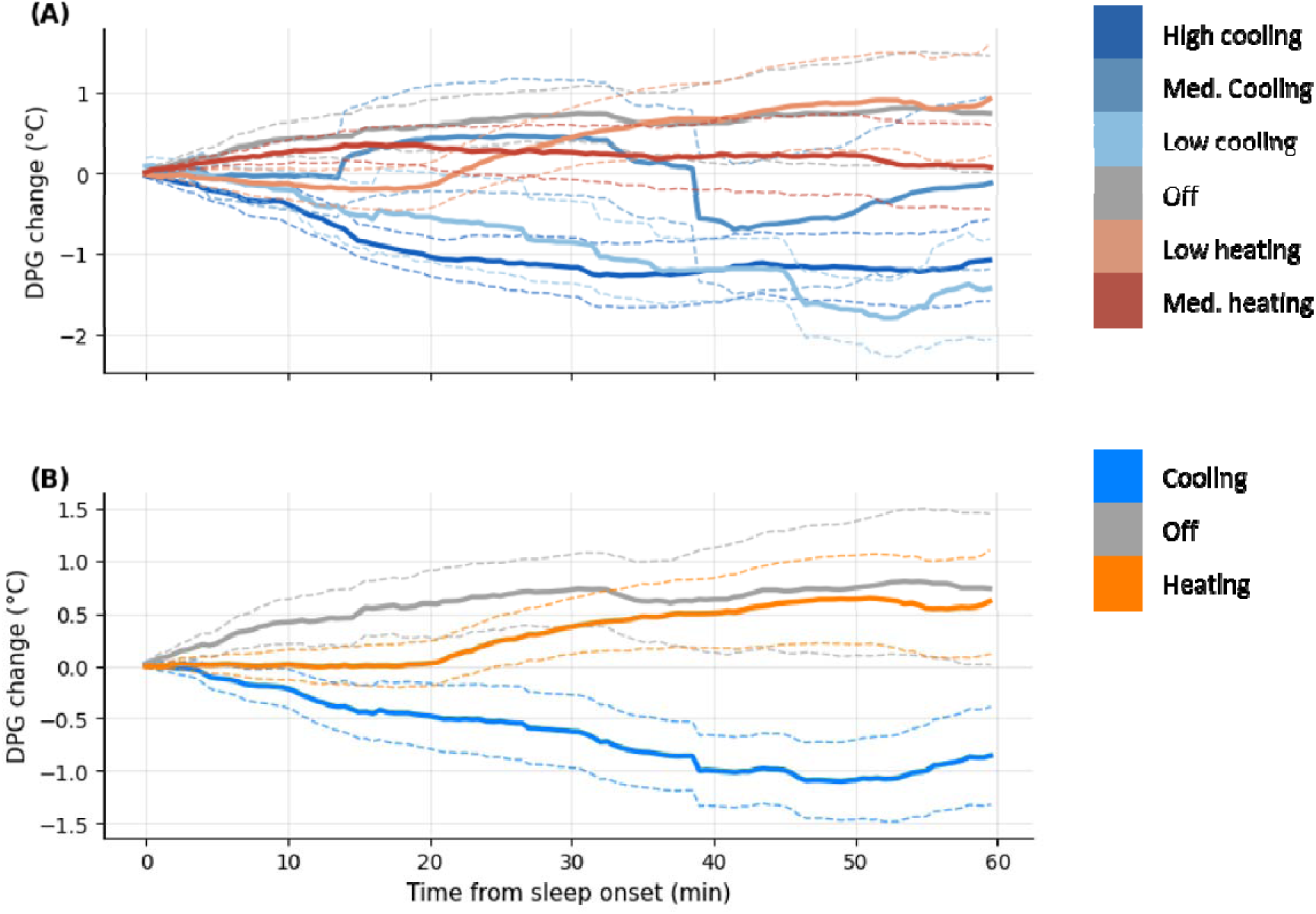
Distal-to-proximal skin temperature gradient dynamics from sleep onset as a function of the first temperature setting. DPG was computed as distal (foot) minus proximal (sub-clavicular) skin temperature (A) Mean DPG change (°C) for each individual setting; dashed lines represent ±1 standard error. (B) Aggregated into Cooling, Off, Heating groups. Point-wise statistical comparisons between condition pairs are reported in Supplementary Figure S3.

## Discussion

This study demonstrates that active, programmable regulation of the sleep microclimate significantly modulates sleep architecture, autonomic activity, and central nervous system activity during sleep. By systematically varying bed temperature across six settings spanning cooling to heating, we characterized temperature-dependent effects on sleep stage architecture across the full night, the early sleep deepening process, heart rate, and EEG spectral composition. Taken together, the findings reveal that thermal interventions act through multiple physiological pathways, and that the direction and intensity of thermal manipulation produce qualitatively distinct outcomes.

### Sleep architecture

The mixed-effects logistic regression models of sleep stage probability revealed a significant effect of active temperature modulation across the entire night. All five active temperature conditions significantly reduced the probability of N1 sleep and increased the probability of N3 slow-wave sleep, indicating a universal shift away from light and toward deep NREM sleep regardless of the direction of the thermal manipulation. The largest N3 effect was observed under high cooling (OR = 3.65), followed by medium heating (OR = 2.64), suggesting that both extremes of the thermal range studied here can promote slow-wave sleep through potentially different mechanisms.

Wake probability was substantially reduced under four of the five active conditions, with the most pronounced reductions under medium heating (OR = 0.31) and low cooling (OR = 0.35). The exception was high cooling, which produced no significant reduction in wake probability (OR = 0.94, p > 0.05) despite generating the largest N3 increase of any condition.

The interesting effect of high cooling, namely largest gain in slow-wave sleep but no reduction in wakefulness, suggests that high cooling may increase sleep depth within sleep but may also increase the propensity for brief awakenings during shallow asleep. The EEG spectral findings offer a candidate mechanistic account of this dissociation: high cooling suppressed theta, alpha, sigma, and beta power, which may promote the emergence of slow-wave sleep. This contrasts with the heating conditions, where significant delta power increases accompanied the architectural shift toward N3, suggesting that heating and high cooling may both increase N3 scoring but through differentiated cortical mechanisms.

The overall probability of being asleep was significantly increased under medium cooling, low cooling, low heating, and medium heating, but not under high cooling, consistent with a non-linear relationship in which moderate thermal stimulation in either direction promotes sleep maintenance, while the most extreme cooling setting may counteract this benefit.

### Falling asleep: sleep onset latency and early sleep deepening

The accelerated sleep deepening observed under first-segment heating conditions aligns with established thermoregulatory models of sleep onset. Heating drove a sustained increase in distal skin temperature, reaching approximately 1.5°C above baseline by 60 minutes post sleep onset, while cooling conditions attenuated this rise and high cooling drove distal temperature below baseline across much of the monitored period. This pattern closely parallels the natural peripheral warming that accompanies the wake-to-sleep transition (Kräuchi et al., 1999; Kräuchi and Wirz-Justice, 1994), and supports the interpretation that bed heating may facilitate the distal vasodilation required for efficient core-to-periphery heat transfer. The DPG dynamics reinforce this mechanism: heating and Off conditions produced positive gradient shifts across the first 60 minutes post sleep onset, reflecting preferential distal warming relative to proximal skin, whereas cooling drove DPG into progressively negative territory, reaching approximately −1.0°C by 60 minutes.

The delta/beta EEG power ratio, evaluated point-wise from sleep onset, provided neurophysiological support for the view that bed heating accelerates early sleep deepening. Heating produced greater sleep deepening than cooling beginning approximately 30 minutes after sleep onset and persisting throughout the monitoring period, and also exceeded Off during the first 10 minutes and again from approximately 35 to 45 minutes. No point-wise differences were detected between cooling and Off at any epoch, suggesting that cooling does not impair the early sleep deepening trajectory relative to a neutral condition, but that heating actively accelerates it. These findings suggest that the primary benefit of early-night heating lies in deepening and consolidating sleep in the first sleep cycle rather than in shortening the wake-to-sleep transition itself; this is also supported by the null finding on sleep onset latency: there was no significant effect of the initial temperature setting on SOL, either across the six individual settings or when collapsed into cooling, off, and heating groups.

While heating conditions tended toward shorter SOL and cooling toward longer SOL at the descriptive level, these differences were not statistically reliable, likely reflecting the limited power afforded by the sample size and the considerable individual variability in thermal preference and thermoregulatory response.

### Heart rate

The heart rate data revealed distinct effects depending on the timescale and analytical frame of the analysis. Point-wise comparisons immediately following sleep onset showed minimal statistically significant differences between conditions despite interesting descriptive trends, likely reflecting the considerable within-person variability in early post-onset heart rate and the limited statistical power of point-wise uncorrected tests in a sample of 18 participants. In contrast, the stage-adjusted mixed-effects models, which accounted for sleep stage and time of night across the full recording, revealed more orderly effects. Cooling produced graded reductions in heart rate that scaled monotonically with intensity: high cooling reduced overall heart rate by 3.06 bpm, medium cooling by 2.10 bpm, and low cooling by 1.18 bpm, all relative to Off. Low heating produced a small overall reduction (−0.28 bpm), while medium heating produced a robust overall elevation (+1.74 bpm).

Stage-specific analyses revealed that cooling-related reductions were most pronounced during N1 and N2, with the largest absolute effect in N2 under high cooling (−5.15 bpm), and were also significant and consistent across REM sleep (−2 to −3 bpm across all cooling conditions). The response during N3 deviated from the monotonic pattern: high cooling remained significantly associated with a heart rate reduction (−2.92 bpm), medium cooling showed no measurable effect, and low cooling produced an increase (+0.99 bpm), suggesting complex stage-specific thermoregulatory-autonomic coupling during deep sleep. Medium heating, by contrast, produced its largest heart rate elevations during N3 (+3.89 bpm), REM (+1.92 bpm), and wakefulness (+4.47 bpm), while effects in N1 and N2 were small and non-significant. The overall pattern, cooling predominantly reducing heart rate in NREM and REM, heating at higher intensities elevating heart rate preferentially in deeper sleep and wake, is consistent with thermoregulatory modulation of vagal tone, and parallels findings from passive high heat-capacity mattress studies in which facilitated conductive heat dissipation was associated with a mean nocturnal heart rate reduction of 2.36 bpm and increased slow-wave sleep (Herberger et al., 2024). The convergence of effects across passive and active thermal interventions strengthens the implication that promoting heat loss from the body’s core, whether through enhanced conduction or programmed cooling, may downregulate sympathetic-autonomic activity during sleep.

### Methodological considerations: temperature dynamics modeling and the use of aggregated conditions

The physics-informed modeling of temperature dynamics, using exponentially increasing and decreasing functions convolved with setting indicators, provided a more realistic and interpretable characterization of how bed temperature changes actually propagate across the night. Because microclimate temperature responds gradually to each setting transition rather than switching instantaneously, simple block-coded regressors would have misrepresented the true thermal exposure at each epoch. The continuous, physics-informed approach avoided this misspecification and yielded better-calibrated estimates of temperature effects on all outcome measures.

A related methodological consideration concerns the use of aggregated Cooling, Off, and Heating groups in two of the secondary, post-sleep-onset analyses (delta/beta and skin temperature dynamics). This aggregation was used solely to recover statistical power for epoch-wise point-wise testing, and was deliberately not used for the manipulation check or the primary sleep-architecture outcome, both of which are reported only at the level of the five individual settings. However, the aggregation pools an unequal number of intensity levels, three cooling settings against two heating settings, so the Cooling and Heating groups do not represent symmetric ranges of thermal intensity, and the aggregated comparisons may understate the variability of effects within each direction. We therefore treat the aggregated secondary findings as hypothesis-generating descriptions of the broad direction of effect (heating versus cooling) rather than as confirmatory tests.

### Limitations

Several limitations should be acknowledged. First, the sample size (18 participants, 36 overnight sessions) limited power to detect modest effects and prevented a thorough analysis of individual differences in thermal responsivity. Second, although the laboratory setting was necessary for PSG acquisition, it may not fully reflect home sleep conditions, where ambient temperature, bedding, and partner presence vary. Third, first-night laboratory effects may have influenced sleep, although the counterbalanced design should have distributed these effects across temperature conditions. Fourth, the microclimate sensors measured temperature at the mattress surface rather than in the air immediately surrounding the sleeper’s body; the exact thermal environment experienced by the sleeper, which depends on blanket coverage, body position, and contact area, was not directly quantified. Fifth, as discussed above, the aggregation of settings into Cooling and Heating groups for two secondary analyses combined an unequal number of intensity levels (three versus two) and should be interpreted as an exploratory, power-recovery strategy rather than a confirmatory comparison. Finally, the point-wise statistical comparisons used in the secondary post-sleep-onset analyses were not corrected for multiple comparisons, so isolated significant epochs should be interpreted cautiously and in the context of the broader temporal pattern.

### Implications and future directions

These findings have several implications for programmable temperature regulation in consumer sleep technology. For individuals who have difficulty initiating sleep or achieving deeper sleep early in the night, bed heating during the first one to three hours may promote sleep consolidation by enhancing peripheral vasodilation and promoting the increase in delta power and corresponding decrease in beta power.

A potentially useful programming strategy may therefore be to begin the night with heating and then transition to cooling: the initial heating phase could support sleep initiation and early sleep deepening, while subsequent cooling could help maintain sleep by promoting cardiovascular downregulation, including reductions in heart rate.

More broadly, the cardiovascular downregulation associated with cooling, particularly the graded and monotonic reductions in heart rate observed across NREM sleep, suggests possible benefit for individuals with elevated nocturnal sympathetic tone, including those with hypertension, stress-related sleep disturbances, or anxiety disorders. However, the non-linearity of the relationship between thermal intensity and sleep outcomes evident in the high-cooling dissociation, together with the substantial individual variability in thermal responsivity, suggests that fixed temperature programs may not be optimal for every sleeper. Adaptive closed-loop temperature control systems that respond in real time to physiological signals such as heart rate, skin temperature, and EEG-derived indices of sleep depth may therefore offer advantages over preprogrammed schedules. It is also important to note that the present findings apply to a specific ambient temperature range; the relative benefits of heating and cooling may differ in warmer or cooler sleeping environments, warranting further study of context-sensitive programming strategies.

Several questions remain for future work. This study examined discrete temperature settings; systematic exploration of finer gradations, and of the apparent ceiling effect observed for high cooling, could identify optimal thermal ranges for different sleep stages and individual characteristics. Long-term studies are needed to establish whether the observed effects are sustained with repeated exposure or attenuate through habituation. Complementing these real-world findings with theoretical modeling of heat exchange, such as heat equation-based simulations of conductive transfer through the mattress and bedding layers, could provide a more mechanistic account of how specific settings translate into microclimate and skin temperature changes.

Extension of this research to clinical populations, including individuals with insomnia, sleep apnea, restless legs syndrome, or thermoregulatory disturbances associated with menopause, could identify which patient groups stand to benefit most from thermal intervention and what programming strategies are best suited to specific conditions. Direct measurement of core body temperature alongside peripheral skin temperature would permit more precise characterization of the heat redistribution dynamics underlying the observed effects, and neuroimaging approaches could clarify the central circuits mediating thermal modulation of slow-wave activity generation.

