## Supplementary Material for "Active Bed Microclimate Regulation Modulates Sleep Architecture, Autonomic, and Central Nervous System Activity during Sleep"

Supplementary Figure S1. Distal skin temperature (DST) dynamics following sleep onset

DST was evaluated point-wise at 30-second resolution from sleep onset, using the epoch immediately preceding sleep onset as baseline. Point-wise two-sided Mann-Whitney U tests were applied at each epoch to identify periods (uncorrected p < 0.05) of difference between conditions.

When examined across individual temperature settings (panel A), DST trajectories diverged shortly after sleep onset. Heating conditions (low heating and medium heating) showed the most sustained rise, reaching approximately 1–2°C above baseline by 60 minutes. The Off condition followed an intermediate trajectory, rising to approximately 1°C. Cooling conditions showed attenuated DST increases, with high cooling driving DST below baseline across much of the monitored period.

When settings were collapsed into Cooling, Off, and Heating groups (panel B), these differences were pronounced. Heating produced significantly higher DST than cooling beginning approximately 5–10 minutes after sleep onset and persisting across nearly the entire remainder of the monitored period (panel C; Mann-Whitney U, p < 0.05). No point-wise differences with p < 0.05 were observed between heating and Off at any epoch (panel D). Cooling produced significantly lower DST than Off during approximately the first 10 minutes after sleep onset, with an additional brief significant window near 12 minutes (panel E), after which the two conditions did not differ significantly.

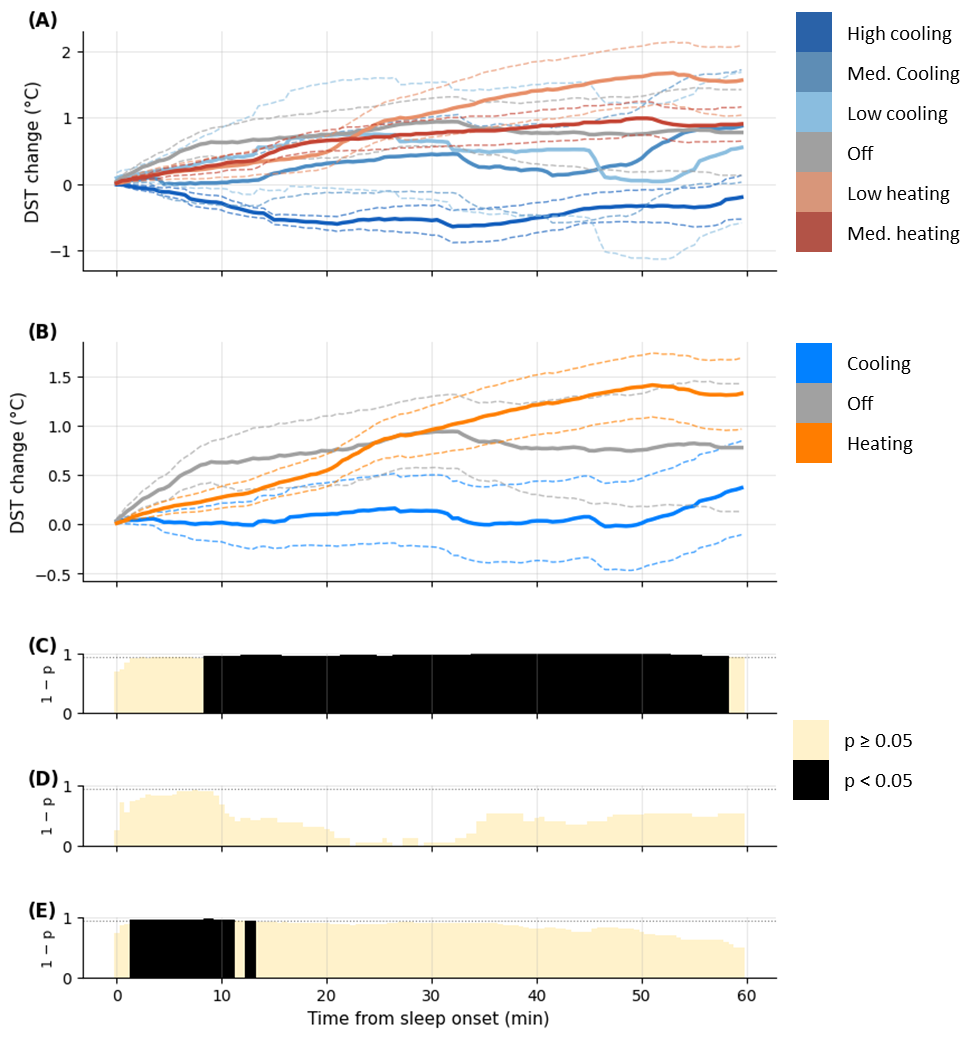

*Figure S1. DST dynamics from sleep onset as a function of the first temperature setting. (A) Individual settings. (B) Aggregated into Cooling, Off, Heating groups. (C–E) Point-wise significance: (C) Cooling vs Heating, (D) Off vs Heating, (E) Cooling vs Off.*

Supplementary Figure S2. Proximal skin temperature (PST) dynamics following sleep onset

PST was evaluated using the same point-wise approach as DST and DPG above. When examined across individual temperature settings (panel A), all active thermal conditions produced a rising PST trajectory across the first 60 minutes following sleep onset, in contrast to the Off condition, which remained largely flat, near 0°C throughout.

When settings were collapsed into Cooling, Off, and Heating groups (panel B), the patterns observed with the individual settings remained consistent. Heating resulted in a brief significantly higher PST in the first few minutes after sleep onset (panel C), followed by non-significant differences for the remainder of the period. Heating produced significantly higher PST than Off across several windows, approximately 20–30 minutes and again from approximately 40–55 minutes after sleep onset (panel D). Cooling produced significantly higher PST than Off from approximately 35–40 minutes and again from approximately 50–60 minutes after sleep onset (panel E).

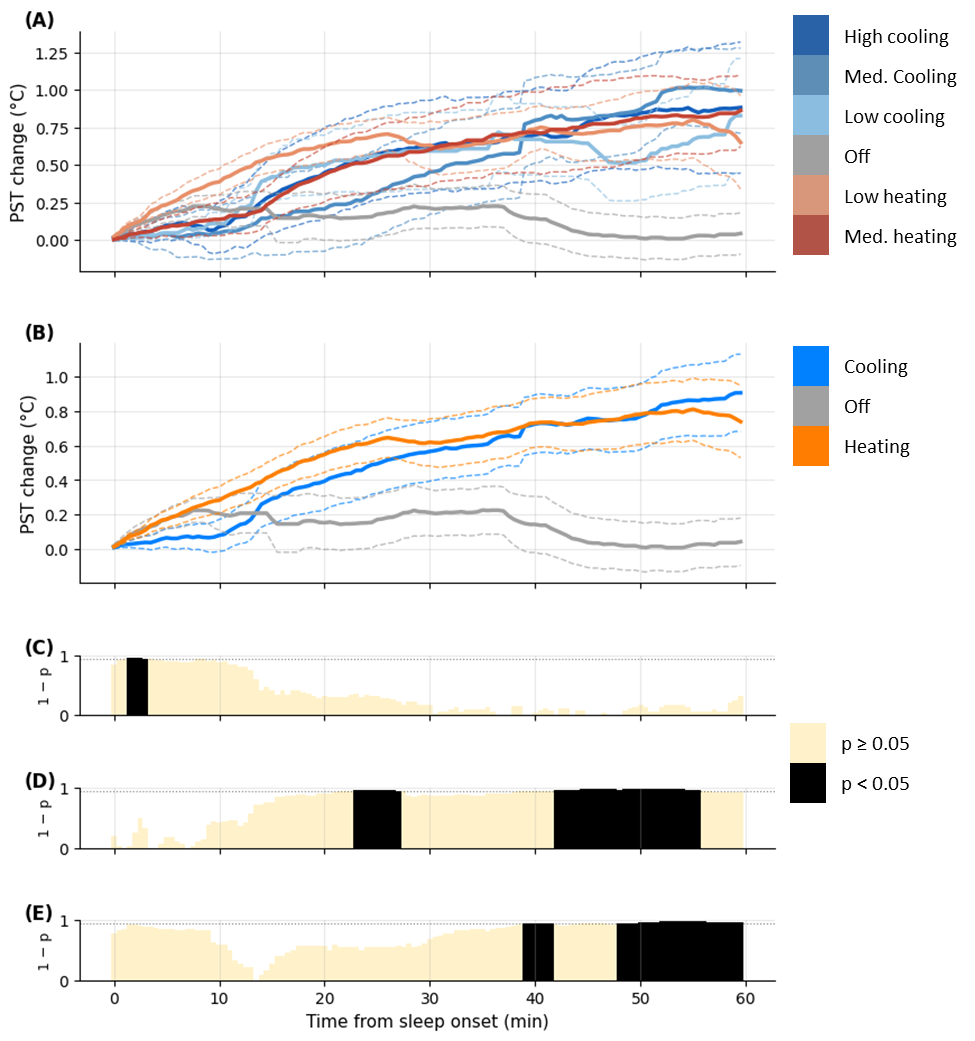

*Figure S2. PST dynamics from sleep onset as a function of the first temperature setting. (A) Individual settings. (B) Aggregated into Cooling, Off, Heating groups. (C–E) Point-wise significance: (C) Cooling vs Heating, (D) Off vs Heating, (E) Cooling vs Off.*

Supplementary Figure S3. Distal-to-proximal gradient (DPG) dynamics following sleep onset

The distal-to-proximal gradient (DPG), calculated as DST minus PST, was evaluated point-wise at 30-second resolution from sleep onset, using the epoch immediately preceding sleep onset as baseline. DPG is presented as the headline skin-temperature secondary outcome because it summarizes the joint distal and proximal dynamics in a single physiologically interpretable measure. Full DST and PST trajectories are reported in Supplementary Figures S1 and S2.

When examined across individual temperature settings (Figure S3), DPG trajectories diverged across the first 60 minutes following sleep onset. Low heating and Off drove a sustained positive DPG shift, rising to approximately +1.0°C above baseline by 60 minutes, reflecting preferential distal warming relative to proximal skin. The medium heating condition produced a mostly flat DPG curve. Cooling conditions drove DPG into increasingly negative territory: high and low cooling produced the most pronounced negative shifts, falling to approximately −1°C.

When settings were collapsed into Cooling, Off, and Heating groups (Figure S3B), heating and Off followed broadly similar positive trajectories, while cooling diverged strongly in the negative direction, reaching approximately −1.0°C by 60 minutes. Cooling produced significantly more negative DPG values than heating from approximately 30 minutes after sleep onset through the remainder of the monitored period (Figure S3C; Mann-Whitney U, p < 0.05). No point-wise differences with p < 0.05 were observed between Off and heating at any epoch (Figure S3D; Mann-Whitney U, p ≥ 0.05 throughout). Cooling produced significantly lower DPG than Off across much of the monitored period, with significant epochs spanning approximately 5 to 20 minutes, and again from 30 minutes onward to the end of the monitored period (Figure S3E; Mann-Whitney U, p < 0.05).

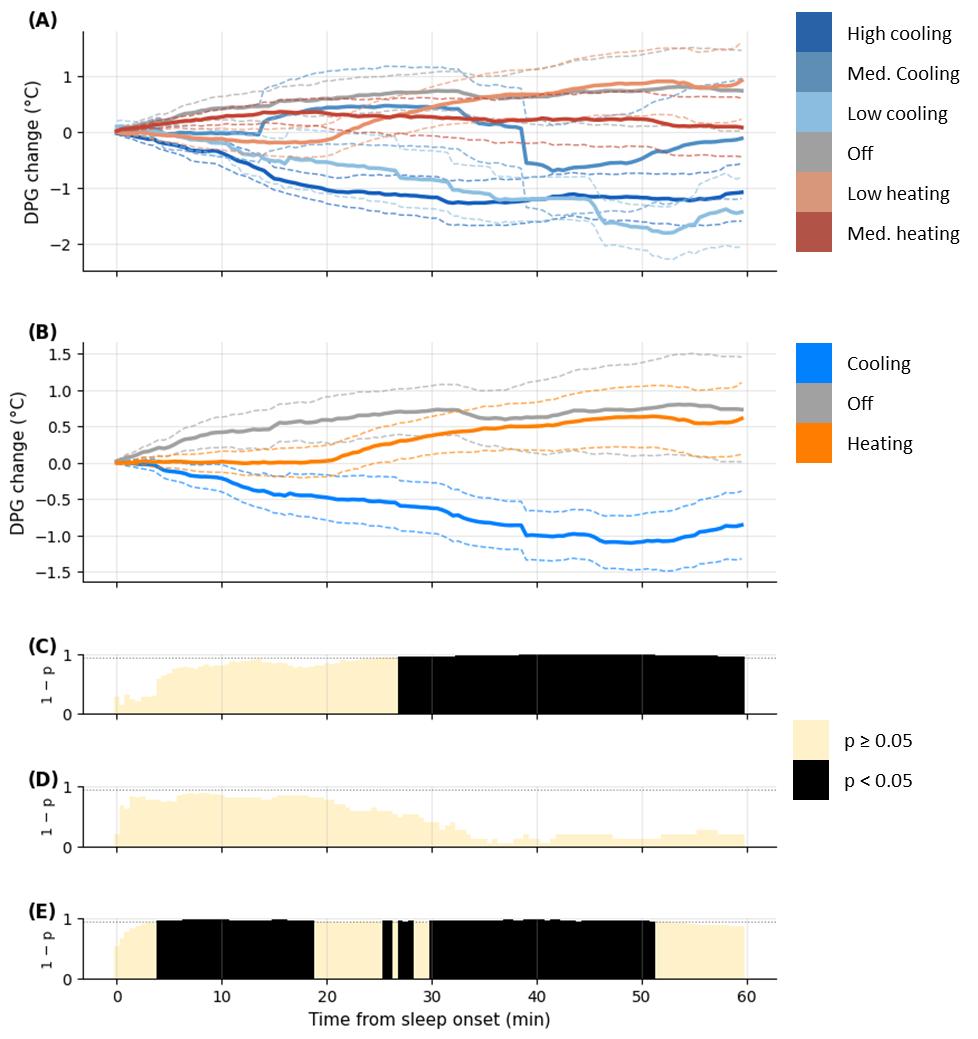

*Figure S3. Distal-to-proximal skin temperature gradient (DPG) dynamics from sleep onset as a function of the first temperature setting. DPG was computed as distal (foot) minus proximal (sub-clavicular) skin temperature. (A) Mean DPG change (°C) for each individual setting; dashed lines represent ±1 standard error. (B) Aggregated into Cooling, Off, Heating groups. (C–E) Point-wise statistical significance (1 − p) from two-sided Mann-Whitney U tests at 30-second resolution. Black bars: p < 0.05; tan bars: p ≥ 0.05. (C) Cooling vs Heating. (D) Off vs Heating. (E) Cooling vs Off.*

Supplementary Figure S4. Heart rate dynamics following sleep onset (point-wise)

Heart rate was evaluated point-wise at 30-second resolution from sleep onset, using the epoch immediately preceding sleep onset as baseline. When examined across individual temperature settings (panel A), heart rate trajectories showed a general tendency to increase above baseline across all conditions over the 60-minute period, with cooling conditions (particularly high cooling) showing the most pronounced rise, reaching 5 bpm above baseline by 60 minutes. Heating conditions remained closer to baseline throughout.

When settings were collapsed into Cooling, Off, and Heating groups (panel B), cooling showed a sustained upward drift reaching > 4 bpm above baseline by 60 minutes, while heating remained near 0–2 bpm across the monitored period. Despite these descriptive differences, point-wise statistical comparisons revealed minimal significant differences between conditions: cooling produced significantly higher heart rate than heating only at a brief window spanning 25–28 minutes after sleep onset (panel C). No point-wise differences with p < 0.05 were observed between Off and heating (panel D) or between cooling and Off (panel E) at any epoch. This point-wise analysis is superseded in interpretive value by the stage-adjusted, all-night mixed-effects model reported as a main-text secondary outcome (Table 5), which provides better-powered and more orderly estimates of the effect of temperature setting on heart rate.

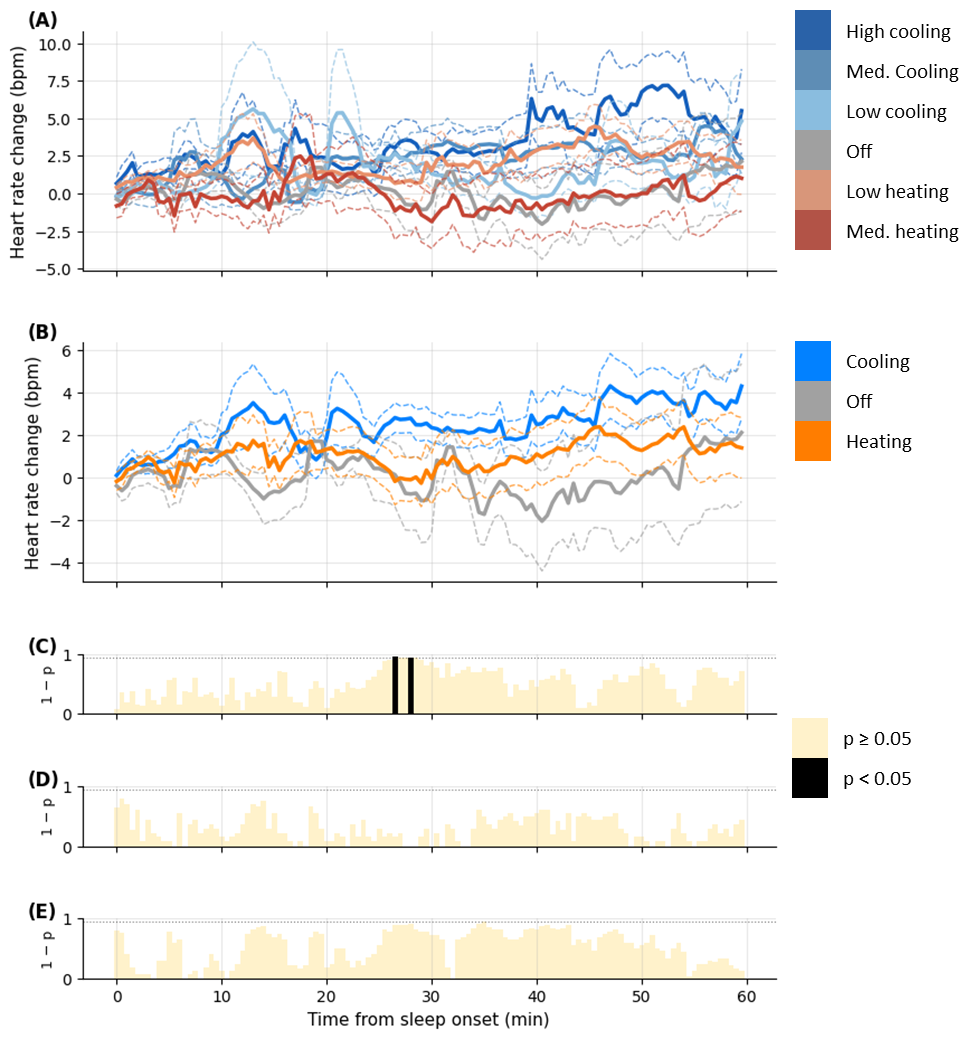

*Figure S3. Heart rate dynamics from sleep onset as a function of the first temperature setting (point-wise analysis). (A) Individual settings. (B) Aggregated into Cooling, Off, Heating groups. (C–E) Point-wise significance: (C) Cooling vs Heating, (D) Off vs Heating, (E) Cooling vs Off.*
